# Fluctuating reproductive isolation and stable ancestry structure in a fine-scaled mosaic of hybridizing *Mimulus* monkeyflowers

**DOI:** 10.1101/2024.09.18.613726

**Authors:** Matthew C. Farnitano, Keith Karoly, Andrea L. Sweigart

## Abstract

Hybridization among taxa impacts a variety of evolutionary processes from adaptation to extinction. We seek to understand both patterns of hybridization across taxa and the evolutionary and ecological forces driving those patterns. To this end, we use whole-genome low-coverage sequencing of 459 wild-grown and 1565 offspring individuals to characterize the structure, stability, and mating dynamics of admixed populations of *Mimulus guttatus* and *Mimulus nasutus* across a decade of sampling. In three streams, admixed genomes are common and a *M. nasutus* organellar haplotype is fixed in *M. guttatus,* but new hybridization events are rare. Admixture is strongly unidirectional, but each stream has a unique distribution of ancestry proportions. In one stream, three distinct cohorts of admixed ancestry are spatially structured at ∼20-50m resolution and stable across years. Mating system provides almost complete isolation of *M. nasutus* from both *M. guttatus* and admixed cohorts, and is a partial barrier between admixed and *M. guttatus* cohorts. Isolation due to phenology is near-complete between *M. guttatus* and *M. nasutus.* Phenological isolation is a strong barrier in some years between admixed and *M. guttatus* cohorts, but a much weaker barrier in other years, providing a potential bridge for gene flow. These fluctuations are associated with differences in water availability across years, supporting a role for climate in mediating the strength of reproductive isolation. Together, mating system and phenology accurately predict fluctuations in assortative mating across years, which we estimate directly using paired maternal and offspring genotypes. Climate-driven fluctuations in reproductive isolation may promote the longer-term stability of a complex mosaic of hybrid ancestry, preventing either complete isolation or complete collapse of species barriers.

**Author Summary:** Hybridization between species can create genetic novelty and promote adaptation, but can also erode species barriers and dilute genetic diversity. Climatic variation likely impacts the extent and eventual outcomes of hybridization, but these impacts are difficult to predict. We use population-scale genomic sequencing of hybridizing *Mimulus* monkeyflowers to better understand the influence of climatic variation on hybridization. We find evidence of hybridization in multiple populations, with groups of different hybrid ancestries clustered along streams in close proximity to each other. Variation in water availability across years appears to affect hybridization between these groups, with less hybridization in drier years compared to wetter years. Paradoxically, this variation may lead to longer-term stability of the hybridization populations, by preventing complete erosion of species barriers while still allowing some gene exchange. In fact, we do see that hybrid ancestry is remarkably stable across a decade of measurements. Climate change is expected to increase the variability of climatic factors such as precipitation and heat events. Our study demonstrates one way these fluctuations could impact species.

## INTRODUCTION

Hybridization can have a wide range of impacts on species, depending on both the success of hybrid individuals and their degree of reproductive isolation from progenitors. At one extreme, unfit hybrids can be an evolutionary dead-end (1–6). Conversely, hybrids may freely and successfully mate with both progenitors, eroding differences between species until a single undifferentiated population remains (7–9). Another possibility is that hybrids are successful but become strongly reproductively isolated from progenitors, forming a new, independent lineage (10–13). Many cases fall between these extremes, with partial but incomplete reproductive barriers (14–17).

Partial reproductive isolation allows for ongoing gene flow between species (introgression) without collapse into a single lineage. Plant biologists have long understood the importance of hybridization among members of a ‘syngameon’, a group of species that exchange genes but maintain distinctiveness (18,19). In the genomic era, introgression has been detected throughout the tree of life, suggesting that partial reproductive isolation is common (20,21).

Introgression plays an important role in numerous evolutionary processes such as adaptation (22,23), niche expansion (24,25), evolutionary rescue (26,27), extinction (28,29), and invasion (30,31). But the dynamics of introgression are varied. Hybridization can be rare and transient, followed by backcrossing within a few generations to leave a signature of introgression at just a few genetic loci (32–34). In other cases, hybridization produces a persistent swarm of intermating hybrids, which then have the potential to interact more extensively with progenitor species (35–38). The coexistence of a partially isolated hybrid swarm alongside progenitor species provides an opportunity for significant adaptive introgression, since hybrids can act as a bridge for gene flow between progenitors (39,40). However, hybridization can also introduce maladaptive alleles or allele combinations (1,41–43) that an admixed population must contend with (37,44–46). Theory suggests that intermediate levels of reproductive isolation may, under certain circumstances, be an evolutionarily stable state rather than simply a transition state on the route to complete speciation (47). But hybridization outcomes appear to be highly contingent on environmental and genetic conditions (35,48–50). More empirical work is needed to understand what conditions lead to persistent partial reproductive isolation in hybrid populations.

One factor that could lead to partial reproductive isolation is spatial heterogeneity. When multiple distinct microhabitats are available, divergent directional selection can maintain multiple ecotypes in parallel niches (51–53). Selection on locally favored alleles in these microhabitats can act as a premating reproductive barrier, reducing gene flow between ecotypes by eliminating migrants (54–57). On the other hand, dispersal between these microhabitats could erode both local adaptation and reproductive isolation between ecotypes, leading to collapse into a single gene pool (7,58,59). Within this pool, multiple microhabitats may instead promote a diversity of alleles through balancing selection (60,61). The relative importance of divergent selection and balancing selection will depend on the scale and frequency of dispersal relative to the scale of environmental heterogeneity (62–65). In some cases, dispersal may be sufficient to prevent complete isolation but not strong enough to erode differences completely. In fact, gene flow is often detected in sympatry even when premating reproductive barriers are strong (66–68). Furthermore, hybrids may establish their own niche in intermediate or underutilized microhabitats, in which persistence is subject to the same balance of migration and selection (11,50,69–71).

Environmental heterogeneity over time, both within and across years, might also contribute to the maintenance of partial reproductive isolation. Differences in the timing of mating throughout the year can isolate groups (72–74). If reproductive phenology is dependent on environmental cues, then variation in those cues across years can modulate the strength of temporal isolation (75–77). Also, variation in selective pressures across years can lead to fluctuating selection that maintains a diversity of genotypes (78–81). Within a hybridizing population, this can result in selection against hybridization in some years but not in others (82,83).

Understanding and predicting the effects of environmental variation on hybridization and reproductive isolation requires tracking hybrid populations across time. Multi-year studies have given us important insights into hybridization, demonstrating directional shifts in composition over time (84), shifts across space (85), stability over time (86,87), and fluctuations in the strength of species barriers (9,82,88). Still, we know little about the environmental and genetic circumstances driving these different outcomes.

With this in mind, we turn to a previously identified hybridizing population of *Mimulus guttatus* and *Mimulus nasutus* monkeyflowers, at Catherine Creek (CAC) just north of the Columbia River Gorge in Washington, USA (Figure 1A). *M. nasutus* diverged from an *M. guttatus* progenitor ∼200KYA, expanding to share much of the *M. guttatus* range across the western United States, where the two hybridize in multiple locations of secondary contact (89). At Catherine Creek and the surrounding area, both species occupy a series of ephemeral seeps, where they co-occur at small spatial scales (90). Previously, *M. guttatus* collections in 2012 from Catherine Creek showed levels of *M. nasutus* genomic ancestry ranging from near 0 to approximately 50%, indicating a history of hybridization followed by backcrossing to the *M. guttatus* parental population (89,90).

**Figure 1.**
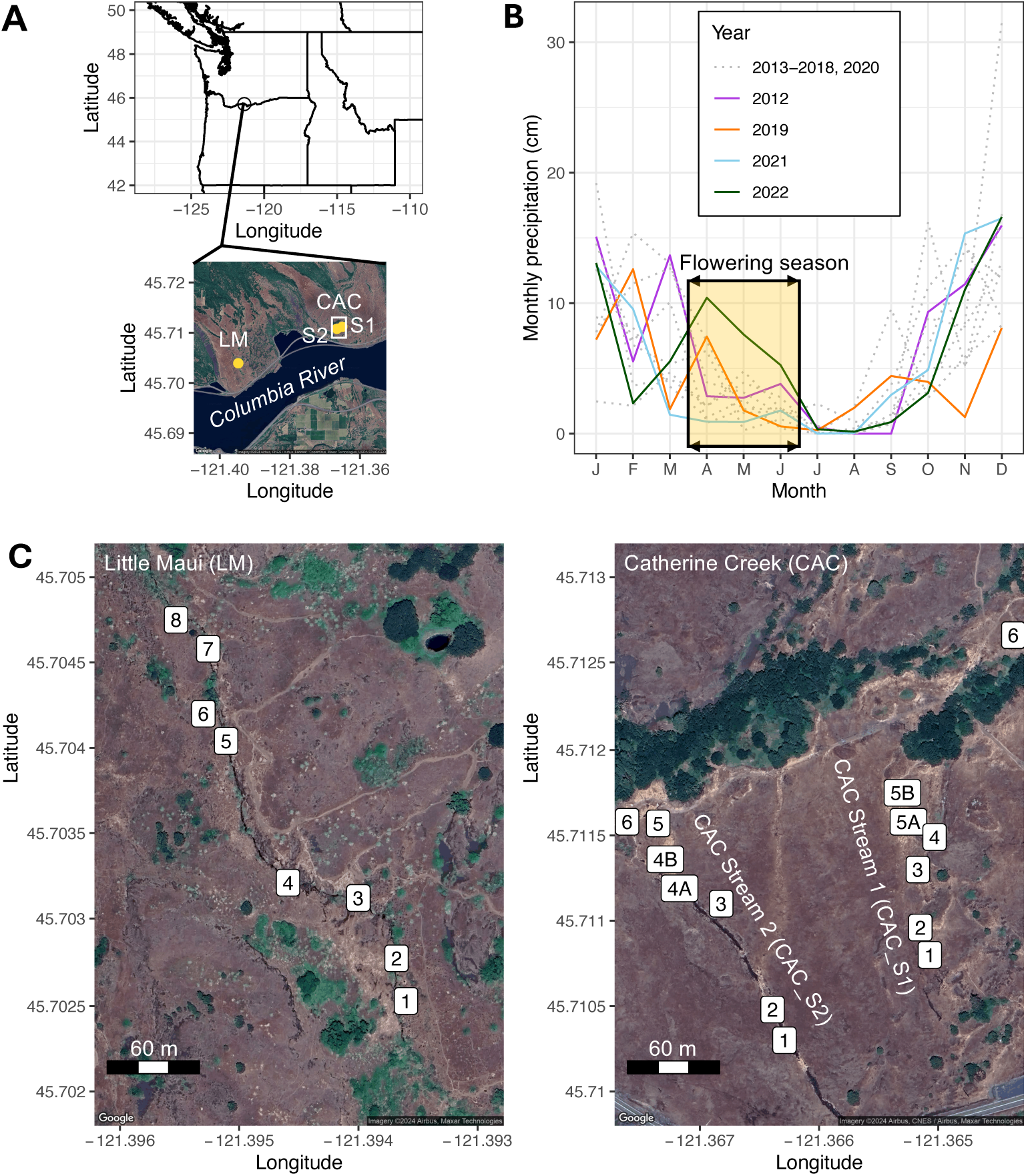
Sampling locations and local variation in precipitation. A) Location of three sampling locations in the Columbia River Gorge area, Washington, USA. LM=Little Maui stream, CAC=Catherine Creek site with two parallel streams, S1 and S2. Approximate distance between CAC_S1 and CAC_S2 is ∼120m, between CAC and LM is ∼4km. B) Interpolated monthly precipitation totals for a 4km grid square covering the CAC and LM sites, from (146). C) Sample plots within each stream. A 0.5mx0.5m square was placed at each plot for flower counts and collections. Exact square placement varied slightly across years, and not all plot locations were sampled in each year.

Despite this evidence of hybridization, several prezygotic reproductive barriers have been documented between *M. guttatus* and *M. nasutus*. One major source of isolation is mating system: *M. guttatus* is primarily outcrossing (though self-compatible), with large showy flowers visited by a variety of bee pollinators (91–93), while *M. nasutus* is predominantly selfing, with small and often cleistogamous flowers that self-pollinate prior to opening (94,95). In addition, the species have phenological differences closely tied to water availability and drought escape: *M. nasutus* tends to be found on mossy rock outcrops that dry out more quickly, while the seepy microhabitats of *M. guttatus* stay wet later into the spring (57,90,96). *M. nasutus* flowers earlier in the season, in part due to a shorter photoperiod requirement for flowering, which has been mapped to multiple large-effect genetic loci (97). In addition, *M. nasutus* is better able to accelerate its life cycle to escape terminal drought (57). The Catherine Creek area has highly variable precipitation (both in quantity and seasonal timing, Figure 1B), so we predict that heterogeneous water availability plays an important role in the persistence and isolation of these species and their hybrids.

Here, we sequence wild-growing individuals and their offspring from three additional years at Catherine Creek and from an additional nearby site, which, combined with previous data, span a decade of collections. This multi-year dataset allows us to ask whether the hybrid population at Catherine Creek is stable across years. By examining fine-scale spatial and temporal patterns of hybrid ancestry, we ask whether environmental heterogeneity might contribute to partial reproductive isolation and hybrid persistence. We address multiple possible contributors to isolation and persistence: spatial segregation of different ancestry cohorts; mating system variation; and phenological isolation across the flowering season. Then, using offspring genotypes, we document how these factors influence the actual mating dynamics of the admixed population, and how these dynamics change across years in response to fluctuating environmental conditions.

## RESULTS

### Introgression from M. nasutus in replicated contact zones

To explore patterns of genomic variation within and between sympatric populations, we collected and sequenced 459 wild *Mimulus* samples from three streams at the Catherine Creek and Little Maui field sites (Figure 1A,C) across the 2019, 2021, and 2022 growing seasons, and performed a PCA on these samples (Figure 2A). PC axis 1 (17.91% of variation) separates species by ancestry, showing a clearly differentiated *M. nasutus* group and a cloud of *M. guttatus-*like individuals with varying levels of hybrid ancestry; PC1 is highly correlated with hybrid index as determined by local ancestry inference (r^2^=0.978, Figure 2B). A separate NGSadmix structure analysis with K=2 is also highly correlated with hybrid index (r^2^=0.979, Figure S1). Additional PC axes reveal population structure associated with geography: PC2 (1.96% of variation) cleanly separates samples collected 4 km apart at the LM and CAC sites (including admixed individuals at PC1 ∼ 0 and *M. nasutus* at PC1 ∼ 0.1), while PC3 (1.74%) and PC4 (1.31%) identify three distinct clusters of admixed individuals within Catherine Creek at a scale of < 300 m (Figure 2A). These three CAC clusters appear to reflect microspatial structure at this site, with two of the three clusters containing only samples from one stream (CAC_S1) and the third containing samples from both CAC streams. Within CAC_S1, each study plot is largely confined to one of these three clusters (Table S3), suggesting genetic differentiation between samples separated by as few as 50 m. Meanwhile, samples from the second CAC stream (CAC_S2) are not differentiated from CAC_S1 samples within their shared cluster.

**Figure 2.**
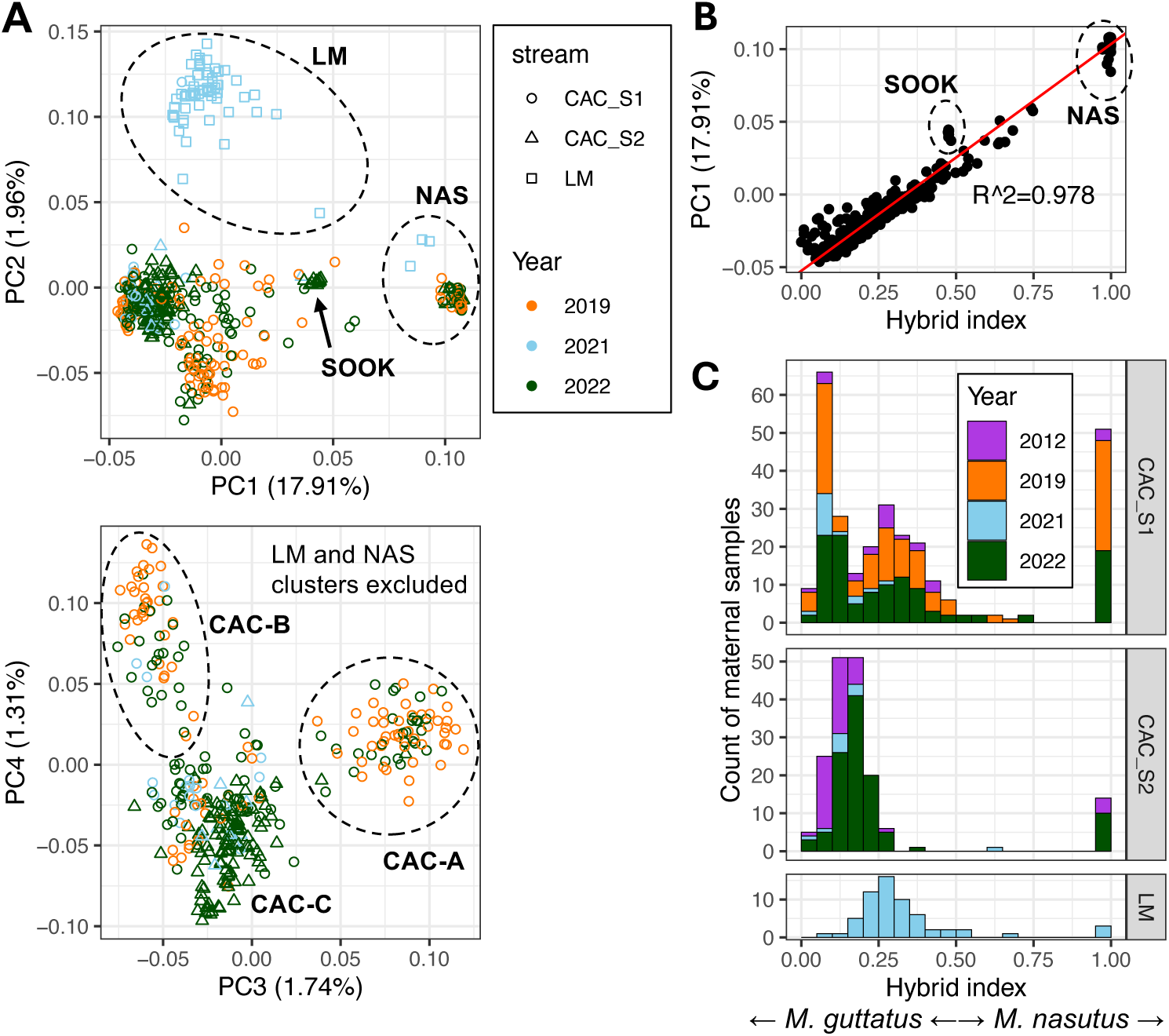
Directional admixture shapes population structure in replicate streams. A) Genomic PCA based on genotype likelihoods at 19,633 variant sites. PC1 separates *M. nasutus* samples (NAS) from *M. guttatus* and admixed samples, while PC2 separates Little Maui (LM) samples from Catherine Creek (CAC) samples. PC3 differentiates an admixed group found primarily in plots S1_1 and S1_2 (cluster CAC-A: note that these plots were not sampled in 2021), while PC4 separates a less-admixed group found primarily in plot S1_4 (cluster CAC-B). The remaining cluster (CAC-C) includes individuals from both CAC streams, which are not differentiated by these PC axes. Variation within Catherine Creek is not structured by year. NAS and LM-A clusters were removed in the PC3-PC4 panel in order to show finer population structure. B) PC axis 1 correlates strongly with hybrid index (the proportion of *M. nasutus* genomic ancestry, determined by local ancestry inference). NAS=*M. nasutus* individuals, SOOK=11 individuals of the allopolyploid species *M. sookensis*. C) Hybrid index, the proportion of sites with *M. nasutus* ancestry across the genome for each individual, is distributed differently in each replicate stream, but this pattern is consistent across years. Hybrid index of 0.0 indicates *M. guttatus*, 1.0 indicates *M. nasutus.* Note that LM was only sampled in 2021; CAC_S2 was only sampled in 2012, 2021, and 2022. Data from 2012 is from (90).

Consistent with previous population genomic analyses (89,90), the distribution of hybrid index (HI, the proportion of the genome with *M. nasutus* ancestry) suggests mostly unidirectional backcrossing of hybrids with *M. guttatus* (i.e., a majority of admixed individuals have HI<0.5, Figure 2C). In fact, virtually all CAC and LM *M. guttatus-*like individuals have at least some detectable *M. nasutus* ancestry: out of 374 majority-*guttatus* samples, only 12 have HI<0.05, while only two of those have HI<0.01. The *M. nasutus* samples, in contrast, have little to no introgression: out of 74 majority-*nasutus* samples, 61 have HI>0.95 and 57 of those have HI>0.99. The remaining 13 have HI between 0.5 and 0.8, indicating that backcrossing with *M. nasutus* does occasionally happen. No individuals were sampled with HI between 0.8 and 0.95, suggesting that backcrossing to *M. nasutus* does not typically continue for multiple generations as it does with *M. guttatus*. Despite pervasive admixture, we detect no first-generation (F1) hybrids (Figure S2). Although we did discover a group of 11 individuals from CAC_S2 with HIs near 0.5, they were determined to be the allopolyploid species *M. sookensis* (Figures 2B, S2, S3, Materials and Methods) and were removed from further analyses of admixture and reproductive isolation. Of the remaining 27 individuals with HIs between 0.4 and 0.6, all have ancestry heterozygosity values less than 0.62 (Figure S2), indicating they are second- or later-generation hybrids as opposed to new F1s (which should have heterozygosity near 1.0). Taken together, our finding that directional introgression composed of later-generation and backcrossed hybrids is replicated in all three sampled streams indicates admixture in secondary contact is not only common but follows consistent patterns across the landscape.

Given the admixture we detect in the nuclear genome, we asked whether organellar genomes also show signatures of introgression. Remarkably, we found that all 325 CAC and LM samples - *M. guttatus,* admixed, and *M. nasutus* - have a single chloroplast haplotype (Figure 3). A similar pattern is seen in the mitochondria (Figure S4). This same haplotype (or close derivatives) is carried by *all* sampled *M. nasutus* and *M. sookensis* lineages, which include collections spanning most of the two species’ ranges (98). In contrast, the haplotype is almost never found in *M. guttatus*, which has much higher organellar diversity across its geographic range (Figures 3, S4): of 18 previously sampled *M. guttatus* accessions, only two carry this CAC/LM haplotype. Surprisingly, one of these two accessions is the only other *M. guttatus* sample in our dataset from a known sympatric site: DPR, which is ∼1000 km south of CAC/LM in the Sierras of California (46,89). We therefore infer that this haplotype is derived from *M. nasutus* and has been completely captured by sympatric *M. guttatus* through introgression at both CAC and LM sites (and, potentially, at other sympatric populations elsewhere in the range). This haplotype structure shows that not a single *M. guttatus* individual in our sample is free from the effects of introgression.

**Figure 3.**
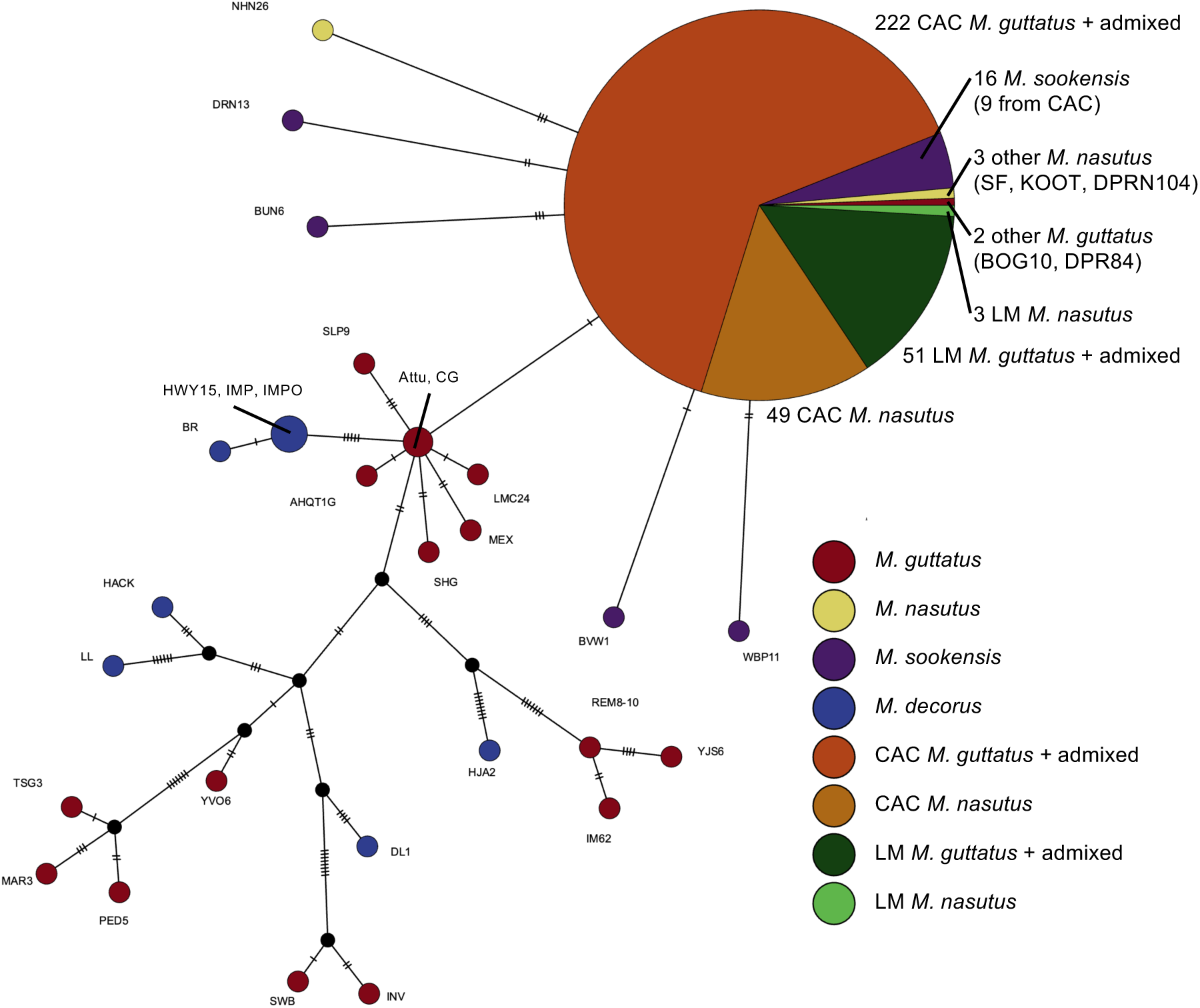
Complete chloroplast capture of a *M. nasutus* haplotype in sympatric *M. guttatus*. NJ-Net haplotype network of chloroplast variation built from 280 CAC maternal samples, 54 LM maternal samples, and 41 additional samples from the *M. guttatus* species complex (98), using 102 total variant sites (43 parsimony-informative). A single chloroplast haplotype is present in all maternal samples from Catherine Creek, including *M. guttatus,* admixed, and *M. nasutus* samples. All *M. nasutus* samples from across the range share this haplotype or a close derivative, as do samples from *M. sookensis* (a polyploid with *M. nasutus* as maternal parent). *M. guttatus* haplotypes are more variable, with only two non-CAC samples sharing the *M. nasutus* haplotype, at least one of which (DPR84) is from another sympatric site with known introgression.

### Patterns of ancestry at small spatial scales are persistent across years

While we find substantial admixture in all three streams, ancestry proportions vary across the landscape (Figure 2C). In each stream, we see multiple distinct, though sometimes overlapping, peaks of ancestry, which we refer to as ancestry cohorts. All three streams have a clear *M. nasutus* cohort (HI>0.95). LM and CAC_S2 each have a second, majority-*M. guttatus* cohort, but with different peak ancestries (HI=0.1-0.2 for LM and 0.25-0.3 for CAC_S2). Within CAC_S1, we see three cohorts: *M. nasutus,* a *M. guttatus*-like cohort centered around HI=0.05-0.1, and a more admixed cohort centered around HI=0.25-0.3. Across multiple years of sampling, the distribution of ancestry in each stream appears similar across years, including the presence of all three cohorts (*M. guttatus,* admixed, and *M. nasutus*) in CAC_S1 and two cohorts (admixed, *M. nasutus*) in CAC_S2. These similarities, despite minor differences in sampling from year to year, suggest that the ancestry structure of these populations is stable across time, even as it varies among streams.

We asked whether the multiple peaks of ancestry within CAC_S1 correspond to spatial structure at a smaller scale. The distribution of hybrid ancestry varies across plots, with clear spatial segregation between the three ancestry cohorts (Figure 4A). These differences are apparent on very fine scales: plots ∼20m apart have distinct distributions of ancestry (i.e., plots S1_3, S1_4, and S1_5A; Figures 1C, 4A). This spatial pattern of ancestry is remarkably consistent across years wherever we have multiple years of sampling. This is true even comparing 2012 (which used a different sequencing methodology) to later years, with a few exceptions: plot S1_5B, for example, appears to have become more admixed between 2012 and 2019. But overall, differences in admixture proportion at small spatial scales are remarkably consistent across a decade of sampling.

**Figure 4.**
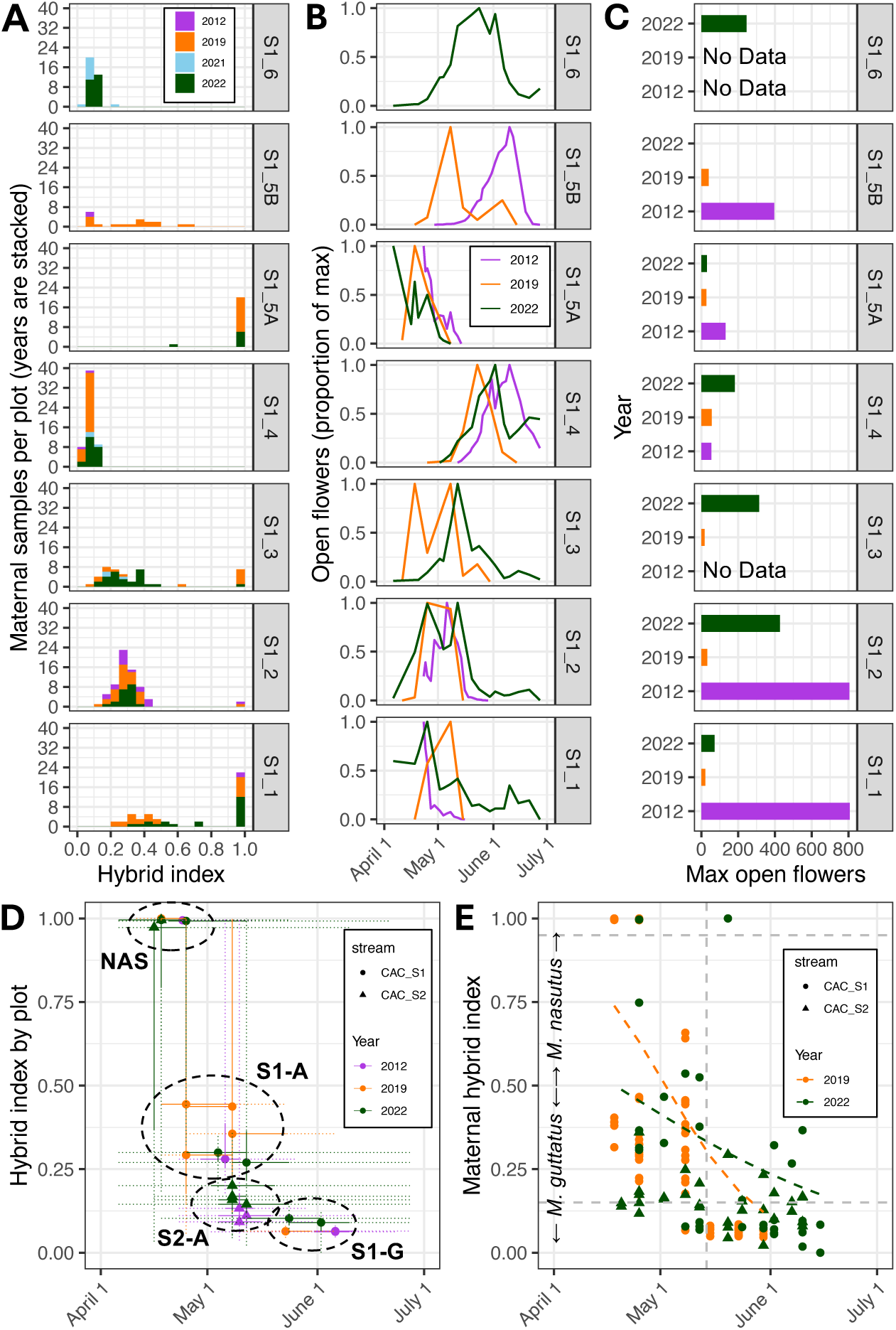
Fine-scaled spatial and phenological structure across years. A) Histogram of maternal samples sequenced from each plot within CAC_S1, binned by hybrid index, with sampling years in stacked bars. Hybrid index of 0.0 indicates *M. guttatus*, 1.0 indicates *M. nasutus.* Seven plots along CAC_S1 vary in the distribution of *M. guttatus,* admixed, and *M. nasutus* individuals, but plots have a consistent distribution across years. B) Counted open flowers throughout the flowering season in seven plots within CAC_1 during the 2012, 2019, and 2022 seasons, scaled to the maximum number of open flowers counted within that plot in that year. Plots with *M. nasutus* individuals (S1 plots 1-3 and 5A) had early flowers, admixed plots (plots S1_1 to S1_3 and S1_5B) had intermediate peak flowering, and plots with *M. guttatus* (plots S1_4 and S1_6) had later peak flowering. Peak flowering was similar but not identical across years. Tails of open flowers indicate that in 2022, admixed plots (plots S1_1 to S1_3 and S1_5B) continued flowering for much longer than in 2019, resulting in more overlap with guttatus plots (S1_4 and S1_6). C) Maximum number of open flowers on any one day for each plot. In 2019, there were far fewer open flowers throughout the growing season than in 2021 or 2022. D) Median flower date for a plot is associated with median hybrid index of that plot. Each point represents the median date out of all counted flowers and the median hybrid index of sequenced individuals from within that plot. Solid lines represent the interquartile range of counted flowers and hybrid indices, while dotted lines represent the full range of flowers and indices for each plot. Circles indicate four main groups: *M. nasutus* plots from both CAC_S1 and CAC_S2, *M. guttatus* plots from CAC_S1, admixed plots from CAC_S1, and admixed plots from CAC_S2. E) Date of marked open flowers is associated with hybrid index of the marked individual. Random sets of flowers were marked when open throughout the season and those individuals were later sampled for sequencing; these are not necessarily the first open flowers for an individual.

### Partial reproductive isolation between admixture cohorts due to mating system

We next sought to investigate potential reasons for the persistence of spatial ancestry structure at such fine scales. One possible cause is premating isolation driven by self-fertilization, so we used maternal and offspring genotype data from 150 maternal families within the two CAC streams to infer selfing rates for our samples. We found that, as expected, *M. nasutus* samples are highly selfing: only 1 of 45 *M. nasutus* offspring was inferred to be outcrossed. *M. guttatus* and admixed cohorts had a mix of selfing and outcrossing, with similar selfing rates between *M. guttatus* (HI<0.15, selfing rates=0.071-0.321, Table 1) and admixed (HI=0.15-0.8, selfing rates=0.218-0.356, Table 1) maternal plants. Among admixed individuals, selfing rate significantly increased with increasing *M. nasutus* ancestry, though this pattern is driven by a small number of individuals with hybrid index >0.5 (p=0.0004 with and p=0.9058 without these individuals, Table S4, Figure S6). Selfing rate also significantly varied across years, with 2022 having the highest selfing rates across ancestry cohorts (Table 1, Table S4, Figure S6). Within the *M. guttatus* and admixed cohorts, selfed offspring were well-distributed across fruits: out of 170 fruits with at least 2 offspring, 51% (86) had a mix of selfed and outcrossed offspring (Table S1). Within mixed fruits, most were majority outcrossing with a mean selfing proportion of 35% (Table S5). Multiple paternity within a fruit was also common: out of 115 fruits with at least two outcrossed offspring, 100 (87%) had more than one inferred pollen donor, with an average probability of 77.6% that two outcrossed offspring in the same fruit were half-siblings rather than full-siblings (Table S5).

**Table 1.** Mating system estimation.

| Cohort* | Year | Maternal |  | Offspring |  |  | Selfing Rate |
| --- | --- | --- | --- | --- | --- | --- | --- |
|  |  | Families | Fruits | Outcrossed | Selfed | Ambiguous |  |
| *M. guttatus*<br>(HI<0.15) | 2019 | 25 | 27 | 112 | 38 | 13 | 0.253 |
|  | 2021 | 15 | 27 | 145 | 11 | 17 | 0.071 |
|  | 2022 | 35 | 35 | 142 | 67 | 6 | 0.321 |
| Admixed<br>(HI 0.15 - 0.8) | 2019 | 26 | 46 | 176 | 49 | 17 | 0.218 |
|  | 2021 | 6 | 12 | 54 | 20 | 5 | 0.27 |
|  | 2022 | 26 | 29 | 123 | 68 | 1 | 0.356 |
| *M. nasutus*<br>(HI>0.95) | 2019 | 8 | 8 | 1 | 24 | 0 | 0.96 |
|  | 2021 | 0 | 0 | - | - | - | - |
|  | 2022 | 3 | 3 | 0 | 20 | 0 | 1 |

\*Determined by maternal hybrid index; includes families from CAC\_S1 and CAC\_S2

We used selfing rates to estimate the strength of reproductive isolation due to mating system. For *M. nasutus*, mating system is an almost complete barrier to reproduction (RI=0.96-1.0, Table 2). Between the *M. guttatus* and admixed cohorts, mating system was a partial, but important, barrier (RI=0.209-0.405, depending on the year, Table 2). Note that these estimates do not consider potential secondary effects of selfing on reproductive isolation, such as lower contributions to the outcrossing pollen pool by self-fertilizing flowers.

**Table 2.** Measurements of premating reproductive isolation.

| Year | Direction*<br>(ovule ← pollen) | Mating system<br>isolation | Phenological<br>isolation | Combined<br>reproductive isolation<sup>†</sup> |
| --- | --- | --- | --- | --- |
| 2012 | G ← N | - | 1 | - |
| 2012 | N ← G | - | 1 | - |
| 2012 | G ← A | - | 0.879 | - |
| 2012 | A ← G | - | 0.989 | - |
| 2012 | N ← A | - | -0.51 | - |
| 2012 | A ← N | - | 0.826 | - |
| 2019 | G ← N | 0.262 | 1.000 | 1.000 |
| 2019 | N ← G | 0.960 | 1.000 | 1.000 |
| 2019 | G ← A | 0.262 | 0.983 | 0.987 |
| 2019 | A ← G | 0.209 | 0.980 | 0.984 |
| 2019 | N ← A | 0.960 | 0.435 | 0.977 |
| 2019 | A ← N | 0.209 | 0.747 | 0.802 |
| 2021 | G ← N | 0.068 | - | - |
| 2021 | N ← G | - | - | - |
| 2021 | G ← A | 0.068 | - | - |
| 2021 | A ← G | 0.294 | - | - |
| 2021 | N ← A | - | - | - |
| 2021 | A ← N | 0.294 | - | - |
| 2022 | G ← N | 0.318 | 0.998 | 0.999 |
| 2022 | N ← G | 1.000 | 0.980 | 1.000 |
| 2022 | G ← A | 0.318 | 0.351 | 0.557 |
| 2022 | A ← G | 0.405 | 0.787 | 0.873 |
| 2022 | N ← A | 1.000 | -0.535 | 1.000 |
| 2022 | A ← N | 0.405 | 0.956 | 0.974 |

Values are scaled from -1 (complete disassortative mating) to 1 (complete assortative mating), with 0 indicating random mating.
\*G=*M. guttatus*, A=Admixed, N=*M. nasutus*.
<sup>†</sup> Combined effects of phenological and mating system isolation.

### Partial phenological isolation between ancestry cohorts

Another potential source of isolation between ancestry cohorts is flowering phenology, so we conducted a multi-year census of flowering phenology at CAC to address phenological isolation. For a given plot, peak flowering times were typically consistent across years (Figure 4B), despite differences in overall flower abundance (Figure 4C). As previously shown for the 2012 growing season (90), we found that plot phenology at CAC consistently tracks hybrid ancestry (Figures 4A-B): median flower date of plots is highly correlated with their median hybrid index (pseudo-*r*^2^ = 0.607, Figure 4D and Table S4). The flowering time of individual plants is also associated with ancestry (pseudo-*r^2^*= 0.326, Table S4): *M. nasutus* flowers were typically marked early in the season, admixed individuals mid-season, and late-season *M. guttatus* individuals later in the spring (Figure 4E).

Focusing on three representative plots in CAC_S1 with consistent sampling across years, we calculated the strength of phenological isolation between *M. nasutus* (plot S1_5A), admixed (plot S1_2), and *M. guttatus* (plot S1_4) cohorts. Isolation between *M. nasutus* and *M. guttatus* was complete or nearly so in all years (RI=0.98 to 1.0, Table 2). This finding, along with the absence of any first-generation hybrids among 459 wild-collected CAC and LM samples (Figure S3A), suggests new interspecific crosses are rare. In contrast, phenological isolation between admixed individuals and *M. guttatus* was variable across the three study years: although the two groups were strongly isolated in 2012 (RI=0.88 and 0.99 depending on direction, Table 2) and in 2019 (RI=0.98 and 0.97, Table 2), they were much less so in 2022 (RI=0.44 and 0.78, Table 2).

What might explain the reduction in phenological isolation in 2022? Although median flowering time of each cohort was relatively stable across years, the duration of flowering was not: admixed plots in 2022 flowered later into the season than in 2019 or 2012 (Figure 4B,D). As a result, there was greater phenological overlap between cohorts in 2022 compared to 2019 or 2012 (e.g., plots S1_1 to S1_3 vs. plot S1_4; Figure 4B,D). We see this pattern in the individual-level data as well: in 2019, all marked flowers after May 15 were from the *M. guttatus* cohort, but in 2022, multiple flowers marked after May 15 were from substantially admixed individuals. Differences in absolute abundance likely play an important role as well: 2019 had much lower flower counts throughout the season in all plots compared to both 2012 and 2022 (Figure 4C).

We also counted relatively more flowers in admixed than in *M. guttatus* plots in both 2012 and 2022 (Figure 4C), explaining the asymmetry of reproductive isolation between the *M. guttatus* and admixed cohorts in those years (i.e., higher probability of pollen flow from admixed to *M. guttatus* plots: Table 2).

The strength of phenological isolation between admixed individuals and *M. nasutus* also varied across years, depending on the direction of pollen flow. Admixed individuals were less likely to receive pollen from *M. nasutus* in 2022 (RI = 0.96) than in 2012 (RI = 0.83) or 2019 (RI=0.65), in part due to a second wave of late-flowering hybrids in 2022 (Figure 4B) that did not overlap with *M. nasutus*. In all three years, *M. nasutus* was poorly isolated from admixed individuals (RI=-0.51 to 0.48: Table 1), but these estimates are incomplete in 2012 and 2022 because our census of open flowers began after *M. nasutus* had already begun flowering (Figure 4B).

### Measured reproductive barriers accurately predict offspring ancestry shifts

Our paired sets of maternal-offspring genotypes allow us to directly test whether the strong premating barriers we detect at CAC actually translate to observed offspring identities. Any deviation in offspring hybrid indices from maternal values implies incomplete assortative mating (postmating barriers could shift allele frequencies at particular loci but are unlikely to cause systematic shifts in offspring HI). For *M. nasutus*, we observed nearly complete assortative mating due to self-fertilization: at CAC only one of 45 offspring across 11 fruits was inferred as outcrossed by BORICE (Table 1), and even this was an intra-*M. nasutus* outcross. We do find one example at LM where 2 of 7 offspring in a single *M. nasutus* fruit have admixed ancestry (HI=0.687 and 0.627), implying a partially-admixed *M. guttatus*-like pollen parent. In the reciprocal direction, one offspring from a 2022 CAC *M. guttatus* fruit (maternal HI=0.089) and one offspring from a 2021 LM admixed fruit (maternal HI=0.316) had hybrid indices consistent with a *M. nasutus* pollen parent. We can therefore estimate that the rate of *M. nasutus* mating outside its own ancestry cohort is 2/67 or 3.0% maternally, and 2/1501=0.1% paternally.

Next, we asked to what extent mating system differences and phenological isolation shape patterns of assortative mating between hybrids and *M. guttatus* at CAC. We reasoned that assortative mating should be maximized in selfed offspring and, indeed, we observe only a slight deviation in the HIs of offspring inferred as selfed relative to maternal values (regression slope = −0.113±0.031, p=0.005, Figure 5A and Table S5). We attribute this slight relationship to uncertainty in either hybrid index estimation or the inference of selfing vs. outcrossing. For outcrossed offspring (which, by definition, have escaped the effects of mating system isolation), we might expect to observe a stronger deviation between maternal and offspring HIs if assortative mating due to phenological isolation is incomplete. This is precisely what we observe: hybrid indices of outcrossed offspring show a clear deviation from maternal values (regression slope = −0.307±0.024, p<0.0001, Figure 5B and Table S5) with *M. guttatus* parents on average producing more-admixed offspring and admixed parents producing less-admixed offspring.

**Figure 5.**
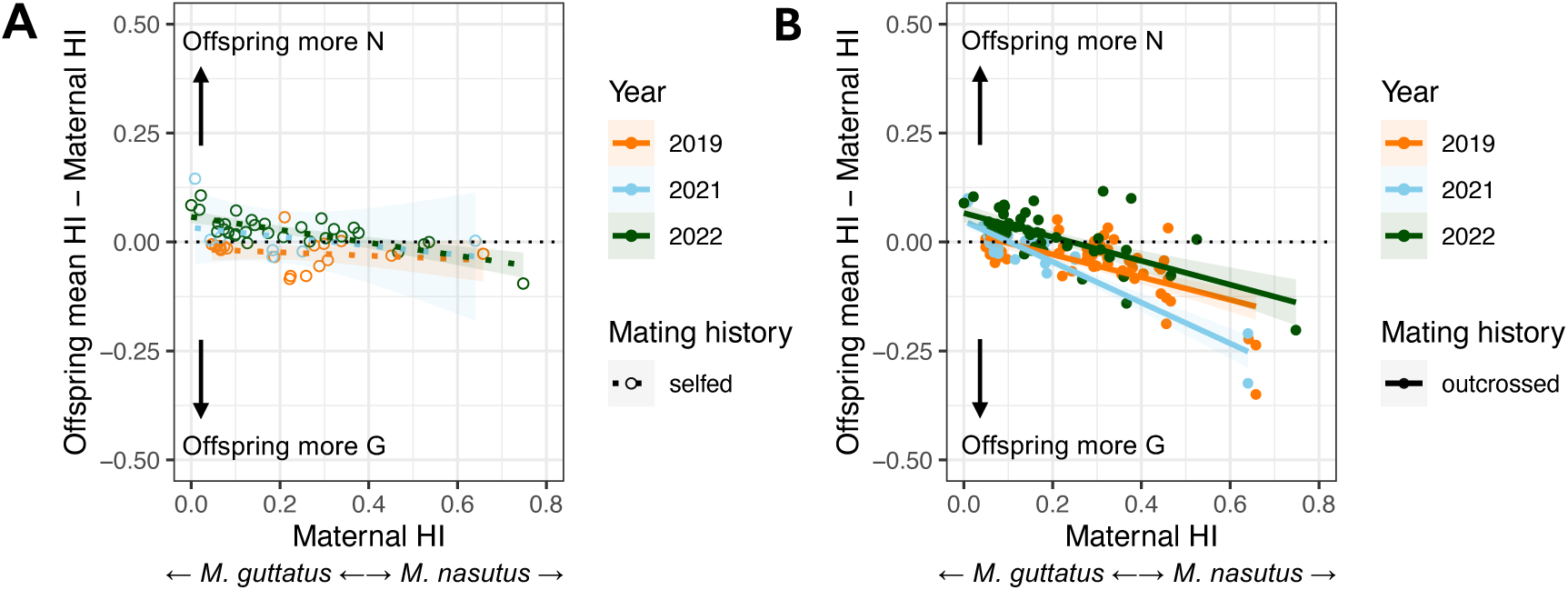
Assortative mating agrees with patterns of reproductive isolation. A) Comparison of offspring hybrid index to maternal hybrid index (HI, the proportion of *M. nasutus* ancestry across the genome) for selfed offspring. Each point represents the mean within a single fruit across all selfed offspring of the difference (Offspring HI – Maternal HI), with positive values indicating greater *M. nasutus* ancestry in offspring relative to maternal plants. Lines indicate individual linear model fits for each year, with shading around each line indicating 95% confidence intervals. The thin black dotted line shows the 1-to-1 null expectation for selfed fruits, where offspring hybrid index is expected to match maternal HI. B) Equivalent plot for outcrossed offspring, showing a larger deviation from maternal ancestry, with the direction of deviation typically in the direction of the population mean.

Strikingly, we also find evidence that yearly variation in assortative mating mirrors patterns of phenological isolation at CAC. In 2019, when phenological isolation between *M. guttatus* and hybrids was nearly complete (RI > 0.98 in both directions), *M. guttatus* maternal plants almost never produced offspring with shifts in ancestry. In contrast, in 2022, when phenological isolation between these groups was at its lowest (RI: 0.34 and 0.79), both *M. guttatus* and admixed parents produced offspring with shifted HIs (Figure 5). In 2019 and 2021, most large shifts in hybrid ancestry occurred in the few admixed individuals with mostly *M. nasutus* ancestry (i.e., HI <0.5), which are further from the population mean; they tended to produce offspring with much more *M. guttatus* ancestry, presumably by crossing with more abundant lower-HI hybrids (Figure 5B). These high-HI individuals were also more likely to self in 2022 (Table 1). The net result of these changes is that, in 2022, outcrossed offspring overall experienced a slight shift to higher levels of admixture compared to maternal samples (mean shift in HI = +0.0172±0.0002), whereas in 2019 and 2021, there was a slight shift to lower levels of admixture (mean shift in HI = −0.0333±0.0002 and −0.0155±0.0004).

## DISCUSSION

Here, we describe in detail the composition and mating dynamics within hybridizing populations of *Mimulus guttatus* and *Mimulus nasutus* during secondary contact. Admixture is prevalent in independent streams, with similar patterns of multi-generational, directional introgression from *M. nasutus* into *M. guttatus*. The distribution of hybrid ancestry across space is variable both at very fine (∼50m) and coarser (∼4km) scales, but stable across a decade of sampling. We measure partial reproductive isolation between three ancestry cohorts by mating system and phenology, both of which likely contribute to the maintenance of ancestry structure across space. We demonstrate substantial year-to-year variation in phenological isolation, which is likely associated with climatic variation. Using direct measurements of offspring ancestry composition, we confirm that variation in measured reproductive isolation predicts observed assortative mating. This is a rare direct confirmation of the effect of premating reproductive barriers on offspring outcomes. Fluctuations in reproductive isolation likely help maintain a mosaic of ancestry across the landscape, preventing either collapse into a single lineage or complete independence of cohorts. Our system demonstrates that the outcomes of hybridization can be dynamic and complex, particularly in scenarios of fluctuating, partial reproductive isolation.

### Repeated cases of introgression across the landscape

While introgression between *M. nasutus* and *M. guttatus* has been detected previously at Catherine Creek and in a separate sympatric area in California (44,46,89,90,99), strong premating reproductive barriers between the two species imply that initial hybridization rates should be rare (66). Our data suggest that ongoing admixture is common across the landscape, with independent admixed populations at Little Maui and Catherine Creek, despite no new hybridization events detected. This follows a trend in other systems where genomic signatures of introgression often co-occur with strong reproductive isolation (68,100), and a broader pattern of frequent signals of hybridization despite generally strong premating reproductive isolation across the tree of life (14,20,21). Theory shows that even occasional migration events can strongly influence allele frequencies (68,101), and rare hybridization events can result in admixed populations if hybrids are persistent once formed. Our finding of weaker reproductive isolation between admixed groups compared to non-admixed progenitors means that, once a few admixed individuals are present, they may promote additional admixture, acting as a genetic ‘bridge’ between otherwise isolated populations and increasing the chance of adaptive introgression (39,40,102,103). Admixed groups might also function as a genetic ‘sieve’: multiple generations of selection can purge incompatible allele combinations, while recombination breaks up linkage between incompatibilities and potentially adaptive alleles (7,46,104).

An open question is how repeatable introgression patterns will be when hybridization occurs multiple times (35,49,104–106). The overall pattern of directional introgression, with pervasive *M. nasutus* ancestry in majority-*M. guttatus* genomes but very little signature of introgression into *M. nasutus*, is consistent across streams. The high selfing rate of *M. nasutus* explains this directionality, which is a common pattern in selfer-outcrosser pairs (77,99,107–109). But despite similar asymmetries, each of our three streams have their own unique distribution of hybrid ancestry. This matches findings in other hybrid zones with a mosaic structure (37,104,106,110,111). Differences in the timing and extent of water availability (77), the spatial distribution of microsites (111), the relative abundance of each species (110), or the presence of pollinators (112) could all influence these idiosyncratic patterns. In the future, expanding this work to additional streams and measuring ecological variables at higher resolution will help us understand which of these or other factors are most important. Overall, we see that a combination of consistent (i.e., selfing) and heterogeneous (i.e., flowering time and microhabitat) reproductive barriers can produce a patchwork of broadly similar but subtly different outcomes each time admixture occurs.

### Complete organellar capture by hybridization

Hybridization often moves organellar genome haplotypes from one lineage into another, a phenomenon known as organellar capture (113–115). Typically, organellar capture is assessed in just a small number of samples at phylogenetic resolution. Our population-scale sample provides a unique window into hybridization history – mainly, that an entire sympatric area has a single maternal origin. An *M. nasutus* maternal origin is consistent with the asymmetric nature of gene flow in our system and the observation, by us and others (66) that *M. nasutus* is more often the maternal parent when hybridizing. It also corroborates our finding that all our *M. guttatus* samples have at least small amounts of nuclear *M. nasutus* ancestry. Still, the extent of capture in even our most *M. guttatus*-like samples is striking and gives us insight into the formation of our hybridizing populations. Not only was the initial hybridization directional, but hybrids must have consistently remained as the seed parent across multiple generations of backcrossing, with nearby *M. guttatus* progenitors contributing primarily through pollen flow rather than seed dispersal. This pattern is consistent with pollen flow acting over longer distances than seed dispersal, so that most new seeds are from maternal plants from within a population, but pollen occasionally arrives from elsewhere. We might expect a similar pattern in other plant systems when pollen flow happens over longer distances than seed dispersal (116), or in animal systems for which males tend to disperse longer distances than females (117).

It is possible that the *M. nasutus* organellar haplotype has some selective advantage within hybrid populations, perhaps due to segregating cytonuclear incompatibilities. Such interactions are common across eukaryotes (118–122), including between populations of *M. guttatus* and *M. nasutus* (123). But we stress that asymmetries in reproductive isolation and dispersal are sufficient to explain these patterns without needing to invoke selection.

### Persistent spatial structure with fluctuating reproductive isolation

Repeated sampling across years allows us to see that each stream has a stable distribution of hybrid ancestry, suggesting a lack of severe hybrid breakdown or maladaptation, although the existence of weak postzygotic barriers (44) helps explain genomic signatures of selection against *M. nasutus* ancestry at Catherine Creek (44,90). Ancestry levels are stable even at ∼20-50m scales within a stream, much smaller than the scale of expected pollinator movement, suggesting that forces other than distance are maintaining isolation between cohorts. Mosaic hybrid zones with stark fine-scaled structure have been described in other systems (124,125) and may be common in plants (126) but there are few studies of change over time in such systems. More broadly, hybrid zone studies have sometimes found stability across years (86,87), and other times found substantial shifts (9,84,85), but the reasons for these patterns are underexplored. We therefore set out to understand what forces might lead to a fine-scaled stable distribution of admixture, and how they might be influenced by a fluctuating environment.

As expected, selfing is an important barrier isolating *M. nasutus* from both *M. guttatus* and admixed cohorts. Interestingly, it also provides a partial barrier between *M. guttatus* and admixed cohorts. For *M. guttatus,* our selfing rates agree with those of other studies, which range from about 25-50% (127–131). Selfing rates in admixed *Mimulus* have not previously been documented; we find that they are generally closer to *M. guttatus* than *M. nasutus*, consistent with dominance of *M. guttatus* floral phenotypes (95). We also see that selfing rates fluctuate slightly across years, possibly due to changes in pollinator abundance and timing (132), or in the size and number of flowers as a consequence of general plant health (133). One intriguing possibility is that the milder conditions in 2022 allowed more plants to have multiple flowers open simultaneously, leading to an increase in geitonogamous (between-flower) selfing, an important selfing mode in *M. guttatus* (129). But overall, selfing is probably a fairly consistent partial barrier across years.

Phenological isolation is also a strong barrier, confirming previous results (90), and is particularly important between the *M. guttatus* and admixed cohorts that are less isolated by mating system. However, phenological isolation is quite variable across years, a result confirmed by differential shifts in offspring ancestry across years. Phenology has a strong genetic component in these species: two major QTL contributing to differences in photoperiod response have been mapped to candidate genes (97). Our results suggest that these genetic differences are only part of the picture, with isolation mediated by other factors. In particular, stronger isolation in drier (i.e., 2012, 2019) compared to wetter (i.e., 2022) seasons points to water availability as a key factor influencing the strength of phenological isolation, acting through changes in flowering duration and abundance. Precipitation amount and variability are frequently associated with shifts in flowering phenology across plant taxa (134–137). With ongoing sampling at Catherine Creek, we will be able to test whether the correlation between precipitation and phenological isolation holds across time. Direct measurements of water availability in microsites across the growing season will also help confirm this relationship in the future.

### Impact of fluctuating environments on hybrid zones

Fluctuating environmental conditions can provide a form of balancing selection, maintaining allelic diversity by favoring different alleles in different years (60,78,80). Similarly, environmental fluctuations may help maintain a variety of ancestry combinations after admixture: (77) found a correlation between interannual variance in precipitation and the extent of introgression across replicate contact zones. Environmental fluctuations could influence the distribution of hybrid ancestry in two main ways: varying the strength of selection against hybrids (83), or directly modulating the strength of premating reproductive isolation (75,76). Our data support the importance of the latter effect, although the former may also play a role. Reproductive barriers that are sensitive to environmental cues, like phenology, may be important sources of fluctuating reproductive isolation across systems.

Climatic variability is projected to increase in many ecosystems with global climate change (138–140). It is therefore imperative that we understand the effects of environmental variability on populations. In the context of climate change, hybridization is predicted to have both beneficial effects, such as increased genetic diversity and adaptive potential, and deleterious effects, such as swamping of rare taxa and homogenization of genetic diversity, each of which is likely to be context- and system-specific (75,141–145). Our study suggests that climatic variability itself can impact the extent of reproductive isolation in hybridizing populations. In a warmer, more unpredictable world, fluctuating reproductive isolation may become more common, leading to an increase in complex, dynamic scenarios of hybridization like this one. To manage these scenarios, we need a better understanding of which factors influence reproductive isolation and hybridization, how they change across space and time, and how they impact the structure and composition of admixed populations.

## MATERIALS AND METHODS

### Environmental data

We obtained daily precipitation data for the years 2010-2022 from the PRISM online database (146) for the 4km grid square centered on (Latitude=45.7113, Longitude=-121.3637), which includes the Catherine Creek site, and took monthly averages.

### Field sampling and collections

We sampled tissue for DNA extraction and sequencing from wild-growing *Mimulus* individuals during the 2019, 2021, and 2022 growing seasons (April through June). In 2019, we sampled from a single stream at Catherine Creek, CAC_S1. In 2021 and 2022, we sampled from both CAC_S1 and an adjacent parallel stream ∼120m to the west, CAC_S2 (Figure 1A). In 2021, we also sampled from a third more distant stream ∼4km away (Little Maui, or LM). In 2021, because we arrived at the field sites in mid-May (due to the ongoing COVID19 pandemic), CAC samples were limited to late-flowering individuals (flowering at LM is shifted to later in the season and thus was unaffected).

Within each stream, we set out 0.5m x 0.5m plots in sites along the stream where *Mimulus* individuals were growing; all samples were collected from within these plots. Plot locations were not always exactly the same across years, but we assigned plots within 10 m of each other to the same plot ID, resulting in a total of 7 plots each for CAC_S1 and CAC_S2, plus 8 plots for LM (Figure 1C); some plot IDs were not represented in every year. Approximately once per week throughout the flowering season in both 2019 and 2022, we counted the number of open *Mimulus* flowers within each plot. The total number of flowers across *M. guttatus, M. nasutus,* and potential hybrids was recorded for each plot at each time point; individual flowers were not field-identified to species as hybrids are difficult to distinguish in the field. Flowers are typically open for 1-3 days, and a single individual may have multiple flowers both simultaneously and in sequence (individual flower number ranges from 1 to >50).

During each visit to CAC in 2019 and 2022 and LM in 2021, we used acrylic paint to mark the calyx of three random open flowers per plot (if available). We used a different color of paint on each visit to indicate the date flowers were open. Later in the season, we attempted to relocate these same individuals and, if successful, collected fruits from the marked flowers, as well as leaf tissue into envelopes with silica for DNA. Seeds from these fruits were later germinated in the UGA Botany greenhouses; leaf/bud tissue was collected from the resulting offspring (N = 1-16 per wild-sampled maternal family, from 1-3 individual fruits per family; details in Table S1) and stored at −80°C for DNA extraction and sequencing.

### DNA Extraction and Illumina sequencing

We extracted genomic DNA from both the dried wild-collected tissue samples and the greenhouse-grown offspring samples using a CTAB extraction protocol with phenol-chloroform extraction (147). Dried tissue was flash-frozen in liquid nitrogen immediately before grinding; fresh tissue from greenhouse-grown offspring was kept at −80C until flash-freezing and grinding. DNA yield was quantified using a Quant-iT DNA quantification kit (Invitrogen P11496) and plate reader, then normalized to equal concentrations for library preparation.

To prepare libraries for Illumina sequencing, we used a Tagmentation approach (148,149); our protocol is available at (150). Briefly, Tn5 enzyme was purified in bulk and pre-loaded with universal Illumina adapters following (151). We then added the loaded Tn5 to approximately 1 ng of sample DNA and incubated to fragment DNA and add universal adapters. Next, we added OneTaq Hot Start polymerase (New England Biolabs M0488L) along with combinatorically barcoded forward and reverse primers and ran 18 cycles of PCR to amplify fragments. After PCR, samples were pooled into sets of 48 and cleaned using SPRI magnetic beads. These sets of 48 samples were quantified with a Qubit fluorometer and then merged in equimolar amounts for Illumina sequencing. Samples were sequenced at the Duke University Center for Genomic and Computational Biology using an Illumina NovaSeq 6000 machine to generate paired-end 150bp reads with a targeted depth of ∼1X coverage per sample. Reads were demultiplexed based on their combinatorial barcodes into individual samples. Samples we sequenced across three separate runs. Across a total of 3055 samples, we sequenced ∼5.5 billion read pairs, for an average of 1.18 million read pairs per sample (standard deviation 2.17 million read pairs). After filtering to remove samples with less than 25,000 called ancestry-informative sites (see *Assigning local ancestry*), our final dataset included 2708 samples with mean sequencing depth of 1.78 million read pairs per sample (standard deviation 1.56 million read pairs, range 45,098 read pairs to 15.72 million read pairs). This final dataset included 459 wild-collected maternal samples and 2248 greenhouse-grown offspring. Details about this final dataset are provided in Table S1.

### Reference alignment, genotyping, and creation of SNP panels

For each Illumina sample, we used Trimmomatic v0.39 (152) to remove adapter sequences and low-quality ends. We aligned reads with bwa v0.7.17 ‘mem’ (153) to the *Mimulus guttatus* IM62 v3 reference genome (https://phytozome-next.jgi.doe.gov), removed duplicates with picard v2.21.6 ‘MarkDuplicates’ (154), and used samtools v1.10 (155) to keep only properly paired reads with map quality >=29. Coverage stats were obtained with qualimap v2.2.1 (156).

We created panels of high-quality, informative SNPs using 36 previously sequenced lines (Table S2), including one *M. nasutus* and 8 admixed *M. guttatus* lines from CAC, plus 3 additional *M. nasutus* and 24 allopatric *M. guttatus* from throughout the species’ ranges. We followed the same steps above to align these Illumina panel lines to the *Mimulus guttatus* IM62 v3 reference genome. The panel was genotyped using GATK 4.4.0.0 HaplotypeCaller and GenotypeGVCFs in all-sites mode (157). Called sites were split into biallelic SNPs and invariant sites, with indels and multiallelic sites removed. SNPs were further filtered with GATK to remove sites with QD<2, QUAL<40, SOR > 3, FS>60, MQ<40, MQRankSum < −12.5, or ReadPosRankSum > 12.5 or < −12.5. Invariant sites were filtered to remove sites with QD<2, SOR>3, or MQ<40. Sites were further filtered at the individual genotype level using vcftools v0.1.16 (158), setting genotypes to missing if DP < 6, DP > 100, or GQ < 15 for that sample. Heterozygous calls were retained. From the resulting genotype file, we created a list of 3,493,514 SNPs called in at least 31 of the 36 reference individuals.

For a more targeted panel, we created a test set of 100 representative wild CAC+LM samples, then subset our variant list to 19,633 sites with nonzero read coverage in at least 60% of test samples and a minor allele frequency of at least 20%. In addition, we created a panel of ancestry-informative sites distinguishing *M. guttatus* and *M. nasutus,* subsetting our full variant list to 208,560 SNPs with >=80% allele frequency difference between the 24 allopatric *M. guttatus* lines and the four high-coverage *M. nasutus* lines.

### Population structure analyses with ANGSD

We used ANGSD v0.940 (159) with the GATK likelihood model to obtain genotype likelihoods for all wild-collected samples at the targeted variant list of 19,633 SNPs. We then used these genotype likelihoods to run a genomic PCA analysis with PCAngsd (160). The resulting covariance matrix was converted into principal components using the eigen function in R (161). As a complementary approach to our ancestry HMM (below), we used NGSadmix (162) with K=2 groups to estimate admixture proportions using the ANGSD genotype likelihoods.

### Assigning local ancestry

We used ancestry_HMM (163) to assign *M. guttatus* vs. *M. nasutus* ancestry across the genome for each maternal and offspring individual, following the *ancestryinfer* pipeline from (164) and using our 208,560 ancestry-informative sites (above). Reads for each maternal and offspring sample were aligned to both the *Mimulus guttatus* IM62 v3 reference and the *Mimulus nasutus* SF v2 reference (https://phytozome-next.jgi.doe.gov), keeping only reads that aligned exactly once to each genome. At each ancestry-informative site, we counted the number of reads supporting each allele. These read counts per allele were used as input for ancestry_HMM (163), along with allele frequencies in the 24 allopatric *M. guttatus* and 4 *M. nasutus* reference lines. Recombination rates between each SNP position were approximated using the bp distance between adjacent sites multiplied by the global recombination rate estimate of 3.9e-8 Morgans/bp, calculated using a total genetic map length of 14.7 Morgans (89) across a genome size of 375 Mb, approximately the mean genome size between *M. guttatus* (∼430 Mb) and *M. nasutus* (∼320 Mb). Ancestry_HMM was run with the model parameters ‘-a 2 0.5 0.5 -p 0 −100 0.5 -p 1 −100 0.5 -e 0.02’ to estimate posterior probabilities of *M. guttatus, M. nasutus,* or heterozygous ancestry at each site for each sample. Posterior probabilities of at least 0.9 for any genotype were converted to hard genotype calls, with lower values set to missing. Samples with fewer than 25,000 called genotypes (out of 208,560 sites) were excluded from all analyses, leaving a final dataset of 2685 samples from an initial set of 3055 samples (Table S1). Hybrid index (HI) equals the number of sites with homozygous *M. nasutus* calls plus half the number of heterozygous calls, divided by the total number of called sites. Ancestry heterozygosity (AH) equals the number of heterozygous calls divided by the total number of called sites.

To test the reliability of our local ancestry measurements, we subset the raw reads from each allopatric *M. guttatus, M. nasutus*, and high-coverage CAC line (36 lines total) to 2 million read pairs per sample, and followed the above local ancestry pipeline to call ancestry across the genomes in these lines. All 4 *M. nasutus* lines had hybrid index >0.99; 18 of 24 allopatric *M. guttatus* lines had HI<0.01 and the remaining six had HI<0.05. The 8 high-coverage CAC *M. guttatus* were more admixed, with hybrid index ranging from 0.11 to 0.34, consistent with previous ancestry estimates for these lines (44).

Our data include 11 maternal samples from plots S2_1 and S2_2, all from 2022, that we determined to be the allopolyploid species *M. sookensis.* These individuals had hybrid indices near 0.5 and ancestry heterozygosity >0.85, which could indicate polyploidy or new F1 hybridization. Offspring of these individuals had very similar hybrid indices and similarly high ancestry heterozygosity, which is consistent with fixed heterozygosity in these highly selfing polyploids (98,165,166) but would not be true for the F2 progeny of F1 hybrids. In addition, these individuals formed a distinct cluster in PCA space: they diverged from the 1-1 correlation between hybrid index and PC1, and they were separated cleanly by PC5. Finally, offspring grown in the greenhouse had a small flower size and shape that matched *M. sookensis* and *M. nasutus;* F1 hybrids typically have larger flowers more similar to *M. guttatus* (Fishman et al. 2002). We removed all *M. sookensis* individuals from the analyses of reproductive isolation, individual phenology, mating system, and offspring ancestry deviations.

We combined our data with previously obtained hybrid indices from 75 individuals collected in 2012 from CAC_S1 and CAC_S2 (90). These samples were genotyped using MSG sequencing (167), and hybrid indices were calculated from an HMM approach implemented in HapMix (168); note that these are different data types and were processed differently than our 2019-2022 data. However, as both approaches use low-coverage whole-genome sequencing and HMM ancestry calling, we expect them to produce broadly comparable hybrid indices even if there are slight differences at individual loci.

### Organellar haplotype networks

To investigate the haplotype structure of maternally transmitted organellar genomes in our population, we aligned each of our maternal samples, along with 8 high-coverage CAC lines (Table S2), to the *M. guttatus* IM767 v1.1 chloroplast assembly (https://phytozome-next.jgi.doe.gov) using BWA v0.7.17 (153), and called genotypes using GATK v4.4.0.0

HaplotypeCaller and GenotypeGVCFs functions (157). Read coverage aligned to the chloroplast genome was high compared to the nuclear genome, so we restricted each analysis to samples with mean coverage >=40, resulting in 328 of our samples and 6 of the previously sequenced CAC lines. We then subset each VCF to a set of SNPs from a recent analysis of organellar haplotypes across *M. guttatus*, *M.nasutus*, *M. decorus*, and *M. sookensis* (98), further removing any SNPs with missing data, heterozygous calls, or individual depth <4 for any sample using BCFtools v1.15.1 (155). We merged this dataset with the genotype calls for the 41 samples from (98), output SNP calls as a fasta sequence for each sample using BCFtools consensus, and converted to nexus format using EMBOSS v6.6.0 (169). We then used the Integer NJ Net function in popart (170) to generate a haplotype network from the resulting alignment. Our final chloroplast dataset included 102 segregating sites, of which 43 were parsimony-informative. We followed this same approach to generate a mitochondrial genome haplotype network, aligning sequences to an *M. guttatus* IM62 mitochondrial assembly (171) and excluding regions annotated as chloroplast-derived. After filtering for mean coverage >=40, we retained 150 of our samples and 8 previously sequenced CAC lines, which were merged with the 41 samples used in the recent analysis. Our final mitochondrial dataset included 120 segregating sites, of which 39 were parsimony-informative.

### Estimating selfing rates

To estimate selfing and outcrossing rates, we first excluded maternal-offspring pairs whose genotypes were incompatible with a maternal-offspring relationship. For each maternal-offspring pair, we used ancestry calls from ancestry_HMM (at 208,560 sites) to calculate the number of informative sites where genotypes were incompatible with maternal-offspring relationships (i.e. maternal and offspring samples were homozygous for opposite ancestries), divided by the total number of homozygous maternal calls with a called genotype in the offspring. Allowing for some uncertainty in ancestry calling, we excluded any offspring for which this incompatibility ratio was >=5%, as well as entire families if greater than half the offspring were excluded. After these filters, our final family dataset contained 185 maternal plants with offspring, plus 1565 offspring.

We then randomly thinned our list of ancestry-informative sites to one per 10kb, leaving 17,514 sites, and calculated genotype likelihoods at each site for both maternal and offspring samples using ANGSD v0.940 with the GATK likelihood model (159). We estimated probabilities of selfing vs. outcrossing for each maternal-offspring pair, and probabilities of half-sibling vs. full-sibling relationships for each outcrossed sibling pair, using the Bayesian model implemented in BORICE.genomic v3 (127). We tuned allele frequencies for 20 steps, then had 20 steps of burn-in and a total chain length of 100, with records kept every 2 steps. Offspring with a posterior probability >= 0.9 were called as selfed or outcrossed, and other offspring were excluded. We counted pairs as half-siblings if the posterior probability was >50%, full-siblings if it was <50%, and missing data if the posterior probability was exactly 50%.

### Calculating the strength of reproductive isolation

We calculated standardized estimates (172) of reproductive isolation (RI) between three ancestry cohorts within CAC_S1, defined by hybrid index: an *M. guttatus* (lower admixture levels <0.15) cohort, an admixed (higher admixture levels >0.15), and an *M. nasutus* cohort. We calculated two components of premating reproductive isolation, mating system isolation and phenological isolation, as well as their combined effects. Each measure ranges from −1 (complete disassortative mating) to 1 (complete assortative mating), with 0 indicating random mating.

For mating system isolation, we used selfing rates estimated from BORICE, dividing individuals into three cohorts by hybrid index. We assumed for this calculation that outcrossing events contribute no isolation to the maternal cohort (RI=0), while selfing events contribute perfect isolation to their maternal cohort (RI=1). The degree of mating system isolation is then equal to the rate of selfing of maternal plants in each cohort. While selfing flowers may also contribute less to the pollen pool, causing increased mating system isolation from the paternal direction, this indirect effect is not included in our calculation.

To calculate phenological isolation, we chose one plot from within CAC_S1 to represent each ancestry cohort: plot S1_4 for *M. guttatus*, plot S1_2 for the admixed cohort, and plot S1_5A for *M. nasutus*. For each pair of plots, we calculated the probability of within-plot vs. between-plot matings based on the number of open flowers in each plot on each day: for each open flower, between-plot mating probability is equal to B/(A+B) and within-plot mating probability is equal to A/(A+B), where A and B are the number of open flowers on the same day as the focal flower from the same and opposing plots, respectively. This assumes that each open flower contributes equally to the pollen pool and ignores the effects of physical distance. These probabilities were then summed across all open flowers across the season to obtain an estimated total number of between-plot and within-plot pairings. Reproductive isolation is then equal to RI=1-2P, where P is the relative probability of these between-plot vs. within-plot pairings.

Following (172), we calculated the combined effect of mating system (RI_M_) and phenological (RI_P_) isolation using Equation 1. Note that the order of isolating barriers does not matter for this calculation.

*Equation 1. Combined Reproductive Isolation*

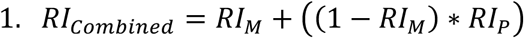

### Using offspring to assess assortative mating

If assortative mating by ancestry is strong, we expect offspring hybrid index to closely match maternal hybrid index; in contrast, deviations between offspring and maternal hybrid index may indicate mating across different ancestry cohorts. To test for deviations from maternal hybrid index within the offspring, we first separated each fruit into selfed and outcrossed offspring based on BORICE. We then calculated the mean for each fruit of the difference between Offspring HI and Maternal HI (Offspring HI deviation), averaging selfed and outcrossed offspring separately. We expect a deviation near 0 for selfed offspring, or under strong assortative mating by ancestry; positive values indicate that the offspring have more *M. nasutus* ancestry than their maternal parent, while negative values indicate they have more *M. guttatus* ancestry than their maternal parent.

### Statistical models

To test for an effect of maternal ancestry and year on selfing rates, we ran binomial GLMs using the R package stats (161). We ran these models for all families, then for all families except *M. nasutus* (HI>0.95) and for only families with maternal HI<0.5.

To test whether phenology was associated with hybrid index at the plot level, we paired flower censuses throughout the growing season from 2012, 2019, and 2022 with hybrid indices from our sequenced maternal plants from those plots. For each plot in each year, we calculated the date that the median censused flower was open in that plot. We then took the median hybrid index of all sequenced individuals from that plot. We ran a beta regression model using the R package ‘betareg’ (173) to test for an effect of median flower date and year on median hybrid index, with year as a factor variable.

We also tested whether flower date was associated with hybrid index at the individual level. For sequenced individuals in 2019 and 2022, we have dates that marked flowers were open – note that these are not necessarily the first flowers on a given individual but were instead randomly chosen flowers from the plot on the day of marking. We ran a beta regression model using the R package ‘betareg’ (173) to test for an effect of individual marked flower date and year on maternal hybrid index, with year as a factor variable.

We ran linear models using the lm function in the R package stats (161) to test for an effect of Maternal HI and Year on Offspring HI Deviation (the difference between offspring hybrid index and maternal hybrid index, averaged per fruit). We ran separate linear models for the selfed offspring and for the outcrossed offspring, as well as a combined model with ‘Mating History’ (selfed or outcrossed) as an additional predictor. We also ran an ANOVA on each linear model using the ‘anova’ function in the R package ‘stats’ (161) to assess the relative effects of each predictor and interaction variable.

For all statistical models, we added each predictor sequentially, allowing for interactions, and used likelihood ratio tests to determine whether the additional predictor significantly improved the model fit.

## ACKNOWLEDGMENTS

We thank Alex Sotola, Sara Hill, Autumn Knight, Liza Lesley, and Sherwin Shirazi for help processing samples, and Bob Schmitz for providing tagmentation enzyme for library preps. Thank you to Alex Sotola, Jill Anderson, Molly Schumer, Casey Bergman, and Kelly Dyer for advice during project development. We also thank Eleanore Ritter, Natalie Gonzalez, Logan Scott, and Sam Mantel for comments on an earlier draft of the manuscript.

## DATA AVAILABILITY

All Illumina data generated for this project will be archived at the Sequence Read Archive (SRA) upon manuscript acceptance. All analysis scripts will be made available on GitHub.

## SUPPLEMENTARY FIGURE CAPTIONS

**Figure S1. Correlation between hybrid index and NGSAdmix structure results.** NGSAdmix value is the proportion assignment to one of two clusters by NGSAdmix with K=2; hybrid index is the proportion of sites with *M. nasutus* ancestry from local ancestry inference using AncestryHMM. The circled ‘SOOK’ cluster that deviates from the 1-1 line is composed of *M. sookensis* polyploid individuals (see Methods). ‘NAS’ = *M. nasutus*.

**Figure S2. Ancestry heterozygosity vs. hybrid index for maternal and offspring plants.** AncestryHMM hybrid index and ancestry heterozygosity outputs for A) maternal and B) offspring individuals. Hybrid index is the proportion of ancestry-informative sites called with *M. nasutus* ancestry from AncestryHMM. Ancestry heterozygosity is the proportion of ancestry-informative sites called has heterozygous (out of the total number of called ancestry-informative sites). First-generation hybrids between 100% *M. guttatus* and 100% *M. nasutus* would have an expected HI=0.5 and AH=1.0; their offspring would be expected to have lower ancestry heterozygosity (∼0.5 if selfed). The circled ‘SOOK’ cluster highlights a group of polyploid *M. sookensis* maternal plants; their offspring have similarly high AH values, which indicates fixed heterozygosity (polyploidy) rather than diploid F1 status.

**Figure S3. PC5 identification of *M. sookensis*.** Principal components 1 and 5 from PCAngsd analysis (PCs 1-4 shown in Figure 2A). PC1 correlates strongly with *M. guttatus* vs. *M. nasutus* ancestry (Figure 2B). PC5 clearly delineates a group of individuals identified as the polyploid species *M. sookensis*.

**Figure S4. Complete mitochondrial capture of a *M. nasutus* haplotype in sympatric *M. guttatus*.** NJ-Net haplotype network of mitochondrial variation built from 147 CAC maternal samples, 11 LM maternal samples, and 41 additional samples (98) from the *M. guttatus* species complex, using 120 total variant sites (39 parsimony-informative). The mitochondrial network largely agrees with the chloroplast network (Figure 3), with a single haplotype present in all maternal samples from Catherine Creek and Little Maui, including *M. guttatus,* admixed, and *M. nasutus* samples. All *M. nasutus* samples from across the range share this haplotype or a close derivative, as do samples from *M. sookensis* (a polyploid with *M. nasutus* as maternal parent). *M. guttatus* haplotypes are more variable, with only one non-CAC-area sample sharing the *M. nasutus* haplotype. *M. decorus* is another member of the *M. guttatus* species complex with variable mitochondrial haplotypes.

**Figure S5. Hybrid index at plot resolution for all three streams.** A) Histogram of maternal samples sequenced from each plot within each stream, binned by hybrid index, with sampling years in stacked bars. Hybrid index of 0.0 indicates *M. guttatus*, 1.0 indicates *M. nasutus*.

**Figure S6. Selfing rate vs. hybrid index.** Proportion of selfed vs. outcrossed offspring per maternal family is slightly correlated with maternal hybrid index (proportion *M. nasutus* ancestry of the maternal plant). *M. nasutus* families are excluded from this plot and the associated model fit (44 of 45 offspring of *M. nasutus* individuals were inferred to be selfed). Self vs. outcross determined by the BORICE Bayesian model. Offspring without >=90% posterior probability of either state were removed. Dashed lines indicate model fits from a logistic regression with formula Maternal HI ∼ Proportion selfed * Year (Table S5).

## SUPPLEMENTARY TABLE CAPTIONS

**Table S1. Summary of maternal and offspring samples by year, stream and cohort.**

**Table S2. Reference panel lines used for SNP panel creation.**

**Table S3. Distribution of samples from each plot in each PCA cluster**

**Table S4. Statistical model output summaries.**

**Table S5. Summary of selfing and sibship estimation from BORICE.**

