## Supplementary figures and images for "Fluctuating reproductive isolation and stable ancestry structure in a fine-scaled mosaic of hybridizing *Mimulus* monkeyflowers"

### Supplementary Figures

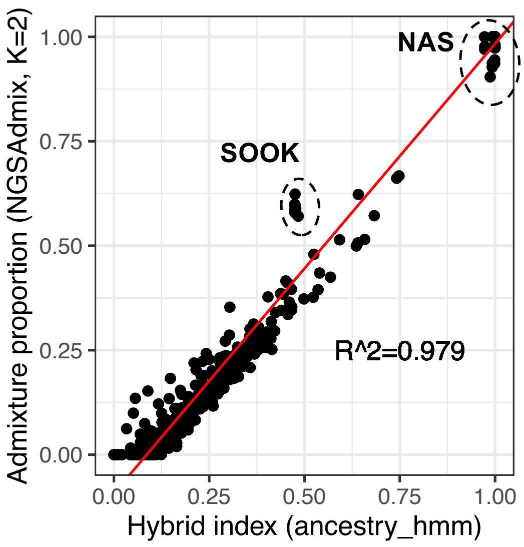

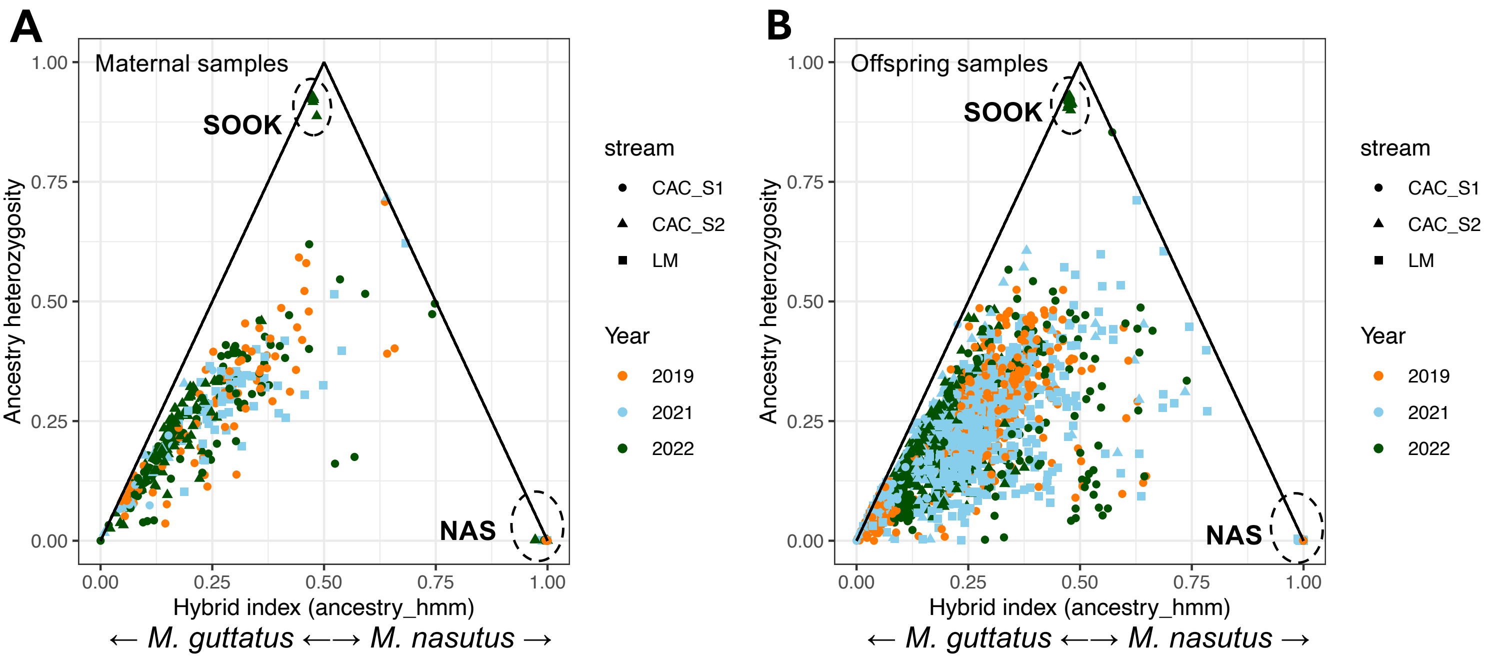

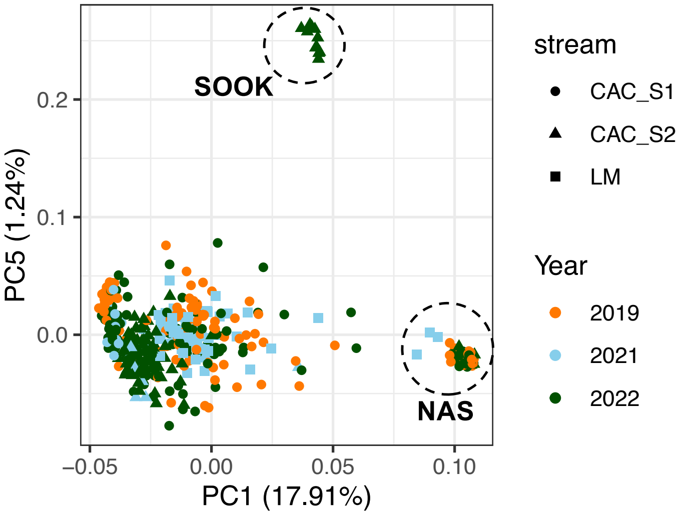

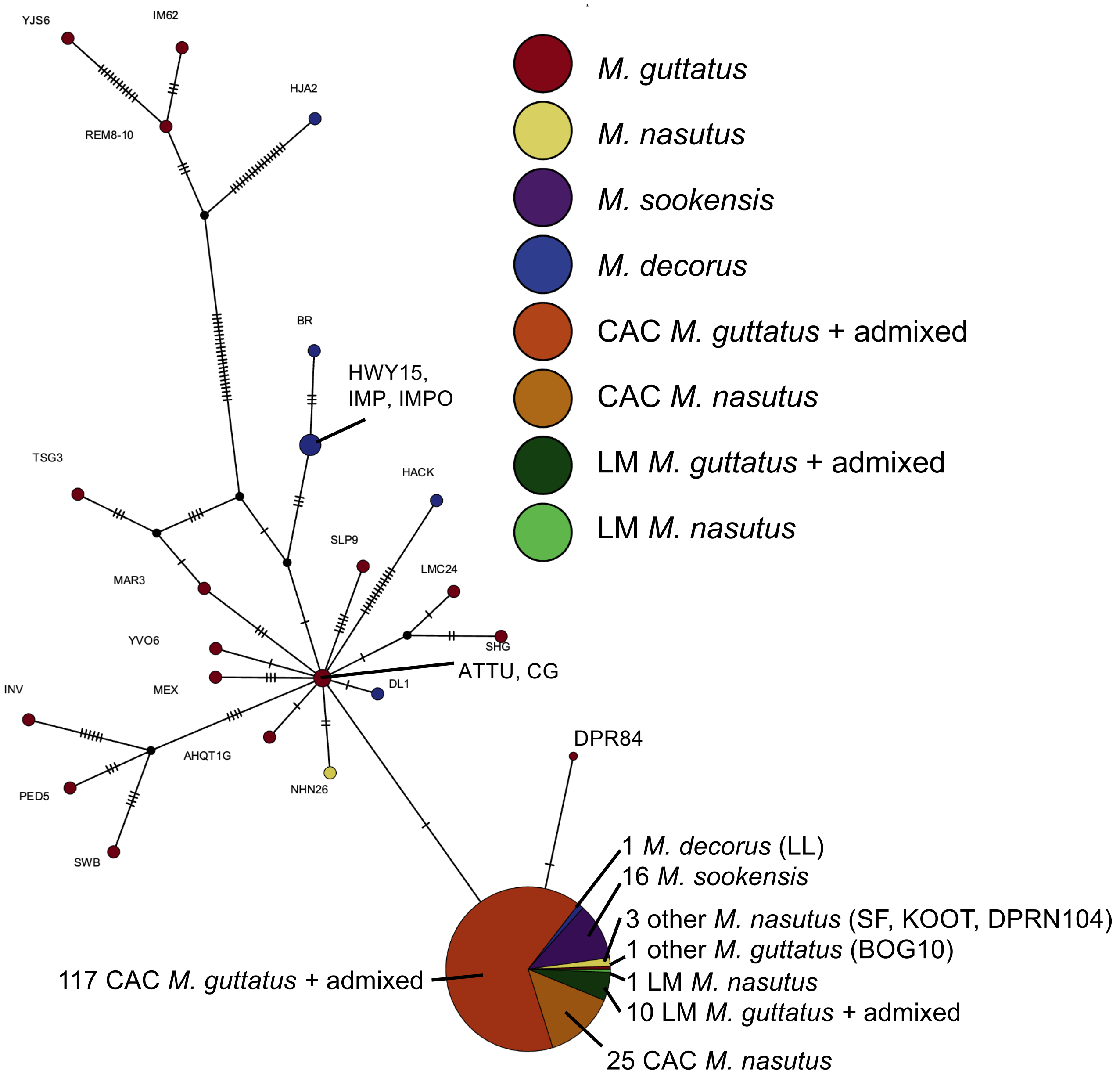

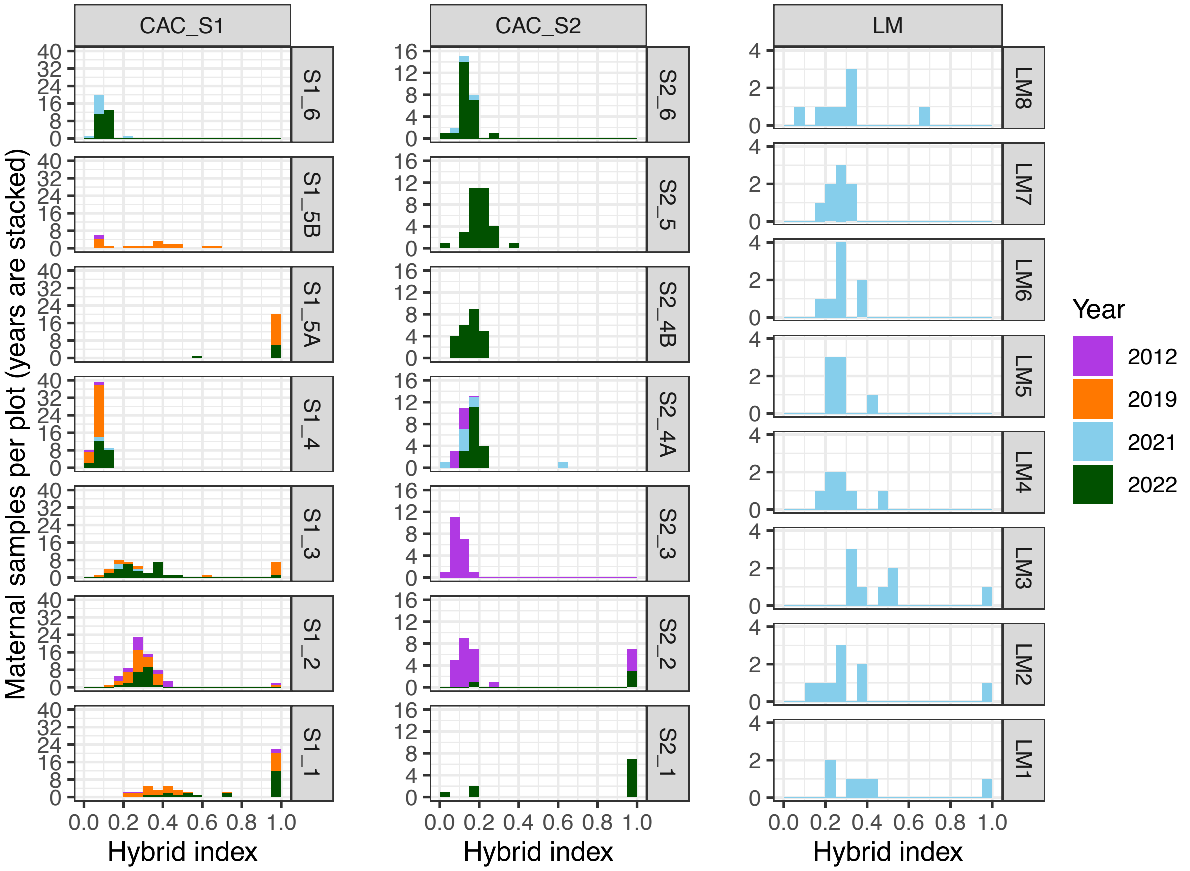

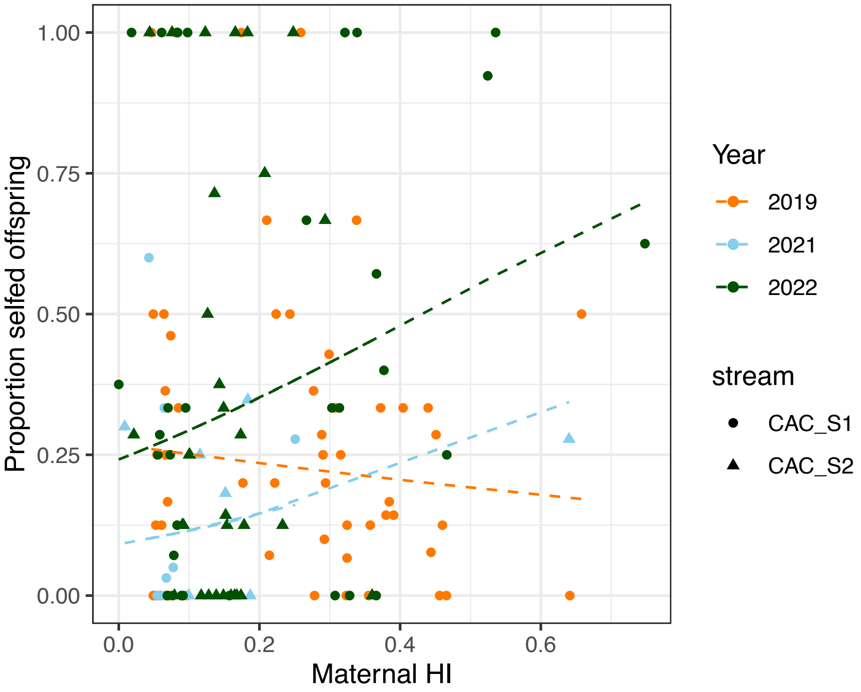
